# Tracheal motile cilia in mice require CAMSAP3 for formation of central microtubule pair and coordinated beating

**DOI:** 10.1101/2021.04.21.440849

**Authors:** Hiroko Saito, Fumiko Matsukawa-Usami, Toshihiko Fujimori, Toshiya Kimura, Takahiro Ide, Takaki Yamamoto, Tatsuo Shibata, Kenta Onoue, Satoko Okayama, Shigenobu Yonemura, Kazuyo Misaki, Yurina Soba, Yasutaka Kakui, Masamitsu Sato, Mika Toya, Masatoshi Takeichi

**Author notes:** **Correspondence:** Masatoshi Takeichi, RIKEN Center for Biosystems Dynamics Research, 2-2-3 Minatojima-Minamimachi, Chuo-ku, Kobe 650-0047, Japan.

## Abstract

Motile cilia of multiciliated epithelial cells undergo synchronized beating to produce fluid flow along the luminal surface of various organs. Each motile cilium consists of an axoneme and a basal body, which are linked by a ‘transition zone’. The axoneme exhibits a characteristic 9+2 microtubule arrangement important for ciliary motion, but how this microtubule system is generated is not yet fully understood. Here we show that CAMSAP3, a protein that can stabilize the minus end of a microtubule, concentrates at multiple sites of the cilium–basal body complex, including the upper region of the transition zone or the axonemal basal plate where the central pair of microtubules (CP) initiates. CAMSAP3 dysfunction resulted in loss of the CP and partial distortion of the basal plate, as well as the failure of multicilia to undergo synchronized beating. These findings suggest that CAMSAP3 plays pivotal roles in the formation or stabilization of the CP by localizing at the basal region of the axoneme, and thereby supports the coordinated motion of multicilia in airway epithelial cells.

## Introduction

Multiciliated cells play a central role in generating directional flow of fluids or particles over the surface of various epithelial tissues, such as oviduct epithelium, ependymal cells lining the brain ventricles, and the airway epithelium of the respiratory tract (Brooks and Wallingford, 2014). Motile cilia on these epithelial cells beat synchronously to produce hydrodynamic forces. Failure of cilia to undergo their coordinated movement causes various diseases, including a subgroup of primary ciliary dyskinesia (PCD) (Lee, 2011; Tilley *et al*., 2015; Damseh *et al*., 2017).

The individual motile cilium of multiciliated cells consists of a 9+2 microtubule arrangement, that is, nine microtubule doublets and a central pair of single microtubules (CP), which position at the outer and central zones, respectively. This core structure of a cilium, called an axoneme, is linked to the basal body (BB), a form of centriole, from which ciliogenesis begins during development (Dawe *et al*., 2007a). The connection between the axoneme and the BB is mediated by a zone called the ‘transition zone (TZ)’ (Czarnecki and Shah, 2012; Goncalves and Pelletier, 2017), and the CP sits just above the TZ, although the outer microtubules extend farther downward to join the triplet microtubules of the BB (Greenan *et al*., 2020). Although extensive studies have been conducted to understand how these ciliary structures form and how motile cilia move in a coordinated fashion (Satir and Christensen, 2007), the molecular mechanisms underlying these questions are still largely unresolved.

CP deficiency brings about defective motion of cilia or flagella (Smith and Yang, 2004; Dawe *et al*., 2007b; Lechtreck *et al*., 2008; Nozawa *et al*., 2013; Loreng and Smith, 2017), and is also associated with PCD (Bautista-Harris *et al*., 2000; Stannard *et al*., 2004; Burgoyne *et al*., 2014). For example, removal of the putative serine-threonine kinase Fused (Wilson *et al*., 2009; Nozawa *et al*., 2013) or radial spoke head proteins such as Rsph1 or Rsph9 (Kott *et al*., 2013; Zhu *et al*., 2019), results in partial or complete loss of CP, simultaneously altering the ciliary beat pattern from planar to rotational in motile cilia. CP formation is thought to occur from TZ. Although molecular constituents of the TZ have been analyzed (Czarnecki and Shah, 2012; Dean *et al*., 2016), how the CP grows out from the TZ in motile cilia, however, remains unknown, particularly because no template for microtubule polymerization is detected at the TZ. In flagella, the CP is anchored to the “basal plate” at the distal end of the TZ. Recent studies identified “basalin’ localized at the basal plate, and its depletion resulted in disruption of the basal plate as well as collapse of CP (Dean *et al*., 2019), suggesting that this structure is important for CP nucleation.

CAMSAP (calmodulin-regulated spectrin-associated protein) and its invertebrate homologs form a protein family, each member of which binds the minus end of non-centrosomal microtubules (Meng *et al*., 2008; Baines *et al*., 2009; Goodwin and Vale, 2010; Tanaka *et al*., 2012; Hendershott and Vale, 2014; Jiang *et al*., 2014; King *et al*., 2014; Richardson *et al*., 2014). Through this property of CAMSAP, it regulates the dynamics of the minus ends of microtubules, simultaneously anchoring them to particular subcellular sites (Nashchekin *et al*., 2016; Toya *et al*., 2016), and contributes to a variety of cell and tissue morphogenesis (Noordstra *et al*., 2016; Martin *et al*., 2018; Takeda *et al*., 2018; Ko *et al*., 2019; Kimura *et al*., 2021; Mitsuhata *et al*., 2021), including neuritegenesis (Chuang *et al*., 2014; Yau *et al*., 2014; Pongrakhananon *et al*., 2018; Feng *et al*., 2019; Wang *et al*., 2019; Chen *et al*., 2020).

The binding of CAMSAP to microtubules is mediated by its C-terminal CKK domain (Meng *et al*., 2008; Baines *et al*., 2009; Hendershott and Vale, 2014; Jiang *et al*., 2014), and its deletion causes the dysfunction of CAMSAP. For example, CAMSAP3, a member of this molecular family, is concentrated at the apical cortex in intestinal epithelial cells of mice through the action of the CC1 domain of the protein, tethering non-centrosomal microtubules to these sites, and this results in the assembly of microtubules whose plus ends point basally (Toya *et al*., 2016). This characteristic microtubule assembly is disrupted in mice bearing a mutant gene encoding a truncated *Camsap3* (*Camsap3*^*dc/dc*^ mice), in which the CKK domain is deleted, because mRNA transcribed from the mutated gene does not cover the exons that encode the CKK domain (Toya *et al*., 2016). *Camsap3*^*dc/dc*^ mice show other defects, such as cyst formation in proximal renal tubules (Mitsuhata *et al*., 2021), and narrowing of the lateral ventricles in the brain (Kimura *et al*., 2021), although the overall brain architecture looks normal. *Camsap3*^*dc/dc*^ mice and *Camsap3* null mutant mice show similar abnormalities (Pongrakhananon *et al*., 2018; Mitsuhata *et al*., 2021), suggesting that CAMSAP3 lacking a CKK domain is fully non-functional.

A recent study showed that CAMSAP3 is expressed in nasal multiciliated cells, and its absence causes defects in ciliary motion and polarity (Robinson *et al*., 2020). Similar observations were also reported using oviduct cells (Usami *et al*., 2021). However, how CAMSAP3 sustains the structure and function of multicilia remains to be investigated. In the present study, we used airway epithelial cells of adult mice to elucidate the role of CAMSAP3 in coordinated movement of multicilia, finding that CAMSAP3 dysfunction due to CKK domain loss results in a partial disruption of axonemal structures at its proximal end, as well as loss of the CP. Superresolution microscopy revealed that a fraction of CAMSAP3 accumulates around the sites where the CP is supposed to be initiated. Based on these observations, we discuss potential mechanisms by which CAMSAP3 supports CP formation and coordinated motion of multicilia.

## Results

### Defective ciliary motion in *Camsap3*-mutated airway cells

*Camsap3*^*dc/dc*^ (*Camsap3* mutant) mice frequently produced clicking or chattering sounds (Audio 1), suggesting that they might have respiratory problems. We examined the luminal surface of trachea collected from postnatal mice using scanning electron-microscopy, finding that multiciliated epithelial cells are normally present in the trachea of the mutant mice, the distribution pattern of which was similar to that observed in wild-type trachea (Figure 1A). In higher magnification views, however, we noted that the alignment of multicilia along the longitudinal axis of trachea, which is generally well ordered in wild-type airway cells, was disorganized to some extent in *Camsap3*-mutated cells (Figure 1B).

**Figure 1.**
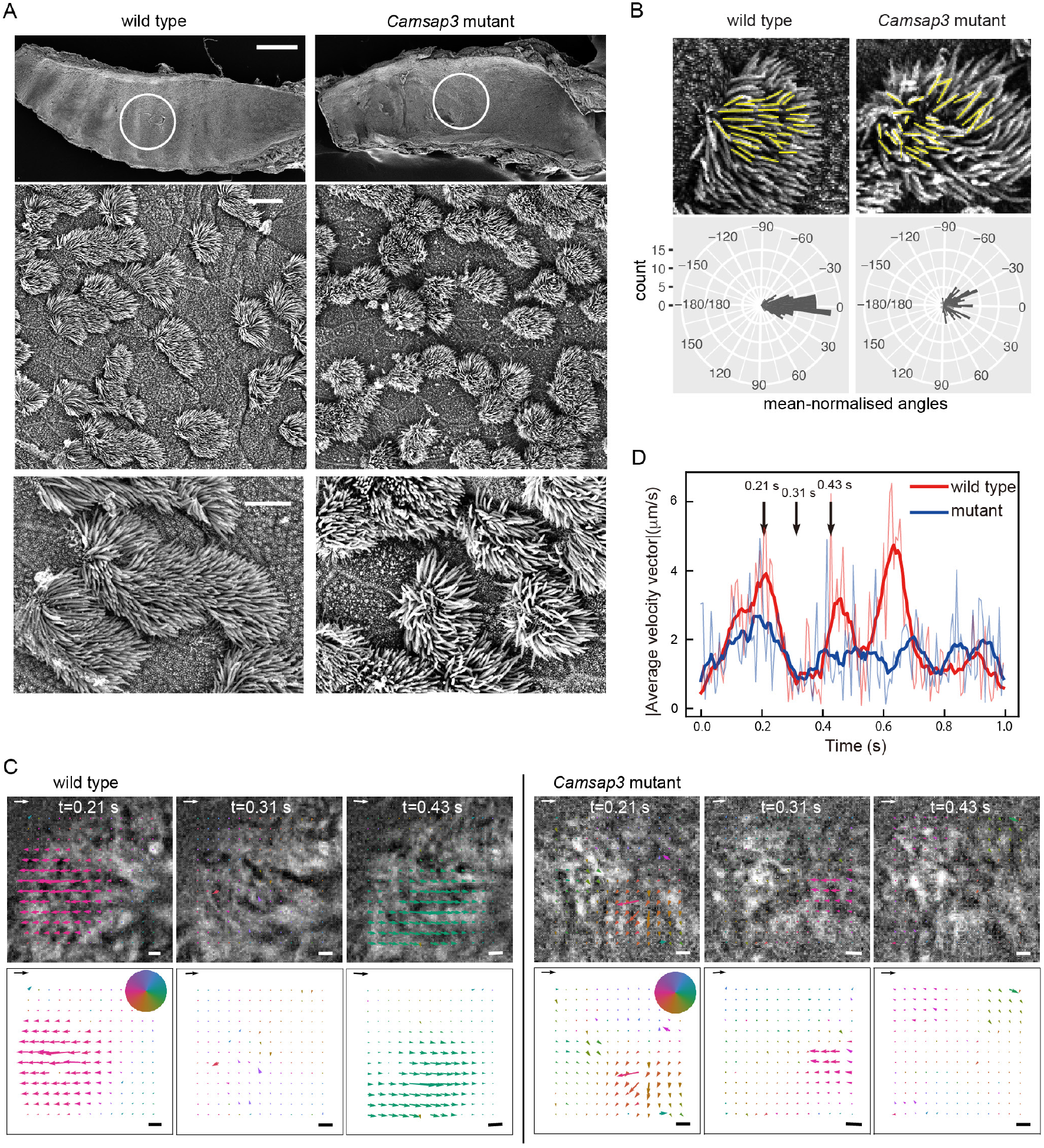
Morphology and movement of motile cilia in airway epithelial cells. **(A)** Scanning electron microscopical observation of the luminal surface of trachea. Trachea was isolated from P21 wild-type or *Camsap3* mutant male mice. The top row shows a whole-mount trachea, in which the luminal surface is exposed. Part of the encircled region was enlarged in the middle and bottom rows. A representative image of three wild-type and five mutant samples is shown. Oral side is at the right. Scale bars: 500 μm, top; 10 μm, middle; and 5 μm, bottom. **(B)** Quantitative analysis of ciliary orientation. Cilia were traced with yellow lines for angle measurement, and polar plots for the distribution of ciliary angles were obtained. The ciliary angles were normalized by the average angle in each cell. Angles of 30 cilia in five cells were measured and aggregated for the polar plots (p < 2.2 × 10^−16^). **(C)** Quantitative analysis of the collective motion of multicilia, which has been recorded in Videos 1 and 2, for a wild-type and *Camsap3*-mutated cell, respectively. These videos were used for PIV analysis (see Videos 3 and 4). Three snapshots, chosen from each video, are overlaid with the flow vector field calculated by PIV analysis, and only the vectors, which represent the velocity vectors, are shown at the bottom row. The color of the vectors indicates the angle of the velocity vector, and the colormap is shown as a pie chart. Scale arrow and scale bar on each image are 20 μm/s and 1 μm, respectively. **(D)** Time-evolution of the magnitude of average velocity vector of ciliary motion plotted for wild-type (red curves) and mutant (blue curves) cells, respectively. The thin curves are the raw data, while the bold curves are the data smoothed by a moving average. The time points shown in B are indicated by black vertical arrows.

To examine whether the motile cilia of the mutant cells have any defects in motility, we isolated trachea and acquired time-lapse images of the cilia on airway epithelial cells. Wild-type cells exhibited a characteristic synchronized beating of multicilia (Video 1), whereas those of mutant cells tended to lose such orderly motion, instead displaying aberrant movements individually (Video 2). We then analyzed the videos to quantify the pattern of ciliary motion by calculating the flow field of a ciliary carpet using particle image velocimetry (PIV). In PIV analysis, we subdivide each image into smaller areas, and then quantify how fast and in which direction each area flows by calculating the cross-correlation between the subdivided areas in subsequent time frames. As a result, velocity is assigned to individual areas, and a flow field is obtained. The flow field for a wild-type cell showed that multiple cilia on the cell oscillated together collectively (Figure 1C and Video 3). In contrast, those in a mutant cell were subdivided into several smaller clusters, each of which showed a collective flow in a unique direction (Figure 1C and Video 4). Using these data, we quantified the degree of synchrony of ciliary motion by calculating the time-evolution of the average speed of the motion, which is the magnitude of the average velocity over the field of view. Since the individual cilia drive oscillatory motion in the flow field of PIV, if all ciliary motions are synchronized, the magnitude of the average velocity vector is expected to show oscillation. In contrast, if the ciliary motion is all spatially random, the magnitude of the average velocity vector would not oscillate in time. Our analysis showed that the magnitude of the average velocity vector oscillated less prominently in the mutant than in the wild-type cells, indicating that the degree of synchrony of ciliary motion was lower in the mutant cells (Figure 1D).

### Loss of the central microtubule pair in the axoneme of *Camsap3*-mutated motile cilia

To investigate the structural basis underlying the uncoordinated beating of cilia in *Camsap3*-mutated cells, we used transmission electron microscopy. Longitudinal sections along the cilia revealed that the overall morphology of cilia and basal bodies looked normal in the mutants. However, the centrally located electron-dense signals that represent the CP were not clearly detectable in the mutant cells, as they were often discontinuous (Figure 2A). Cross sections of cilia revealed further details of such abnormalities. While the cilia in wild-type cells exhibit a typical 9 +2 arrangement of microtubules, any cilium of mutant cells did not have the CP. In these cells, 9 outer doublet microtubules were still present in 70% of cilia at the normal positions, but in the rest of them, one of the doublet microtubules was displaced to an inner zone of the axoneme (Figure 2B).

**Figure 2.**
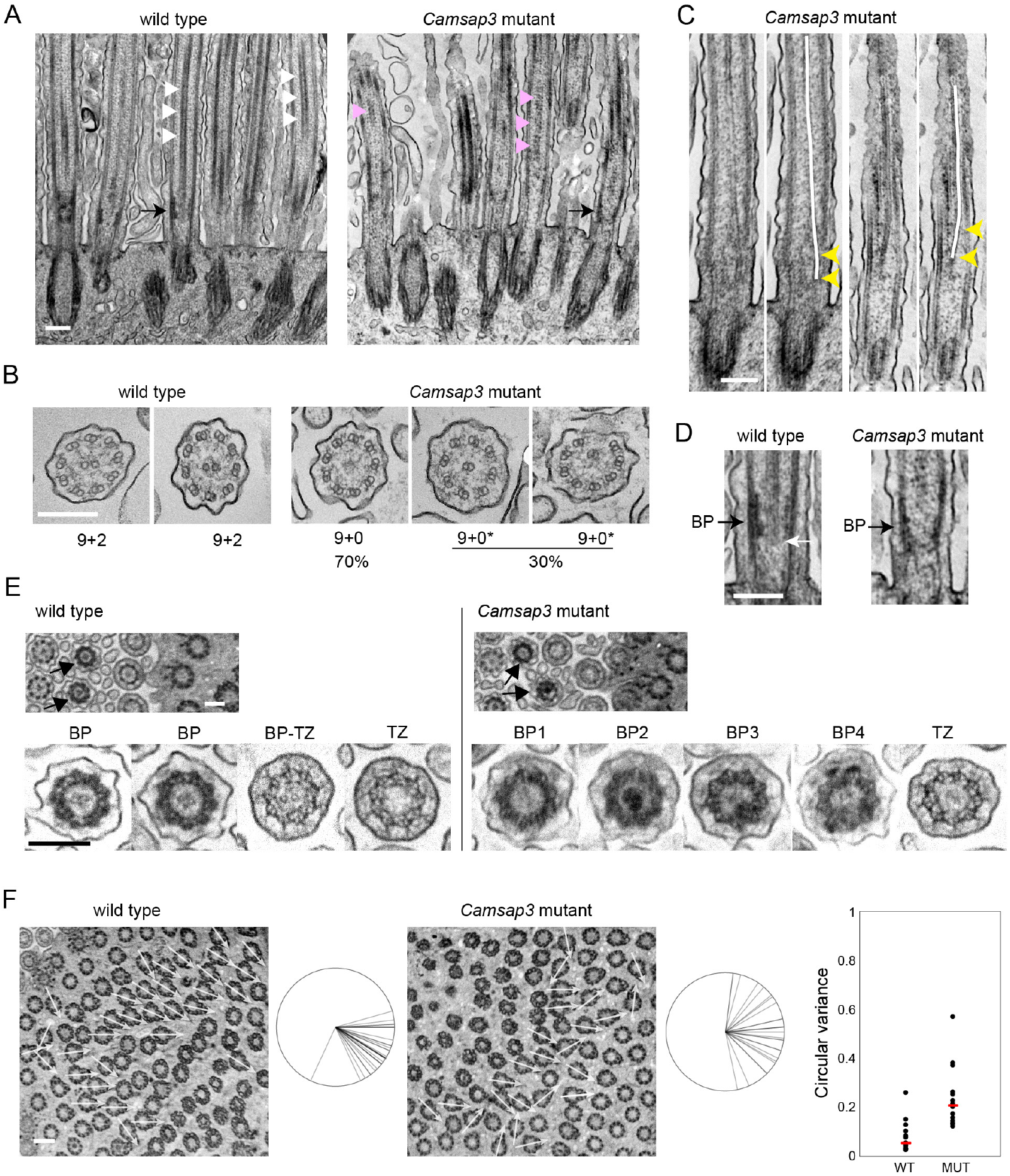
Electron microscopic analysis of cilia and basal bodies. **(A)** Longitudinal section of a multiciliated cell, which yields a longitudinal view of axonemes and basal bodies. White triangles indicate examples of the CP in a wild-type cell; magenta triangles point to intermittent structures at the central position of axoneme in a *Camsap3*-mutated cell. Black arrows point to the ‘basal plate’, and a corresponding position in the *Camsap3* mutant sample. A representative image of more than 50 sections is shown for each genotype. **(B)** Transverse section of cilia. In the mutant samples, 218 cilia from two independent sections were used for analysis. The mutant axonemes did not contain a CP, instead some of them (marked as 9+0*) had a doublet microtubule that was mislocalized to central or semi-central positions. **(C)** Longitudinal section of cilia in *Camsap3*-mutated cells. A centrally located microtubule structure tilts toward the peripheral zone at basal portions of axoneme, as indicated by yellow arrowheads. Two examples are shown, each of which is duplicated to trace the central microtubule with a white line over the image. **(D)** Part of the image in A is enlarged. White arrow points to the proximal end of the CP. BP, basal plate. **(E)** Slightly oblique, transverse sections of a multi-ciliated cell, showing cross-sectional views of axoneme and BB at various longitudinal levels, in which arrows point to BPs that are characterized by a dense peripheral ring. Typical examples of ciliary sections at the level of BP and TZ are enlarged at the bottom. The image labeled BP-TZ likely represents a section at the boundary between BP and TZ. In the mutant sections, all of the BPs show some abnormality, including loss or over-condensation of the central structure, and partial deformation of the peripheral ring. Typical examples are shown. More than 20 sections, each of which contains more than 50 cilia, were examined for each genotype. **(F)** Slightly oblique, transverse section of a multi-ciliated cell, focusing on the basal foot of basal bodies. The direction of individual basal feet is shown with white arrows on the image, and these are aggregated at the right of each image. The range of orientation of basal feet on the cell varied from 26° to 108° in the wild type, with one exceptional case, and 13° to 172° in the mutant sample. We counted 33 and 26 basal bodies for the wild-type and mutant samples, respectively. Graph shows the variation in basal foot polarity in 13 wild-type or 14 mutant cells, which was quantified using circular variance (CV) that was defined previously (Shi *et al*., 2014), where cells with lower CV represent uniform BB orientation. Red bar, median. WT, wild type; MUT, *Camsap3* mutant. Age of mice used, P126. Thickness of sections, 50 to 70 nm. Scale bars, 200 nm in A-E; 400 nm in F.

To seek the origin of these inner doublet microtubules, we reexamined the longitudinal sections and found that, when centrally located microtubules were detectable, they were tilted along the longitudinal axis, one end of which gets closer to peripheral regions of the axoneme around its basal positions (Figure 2C). These observations suggest the possibility that an outer microtubule doublet was abnormally internalized at upper regions of the axoneme, as observed in PCD (Burgoyne *et al*., 2014).

We then focused on the proximal zones of axonemes, where the CP is thought to be nucleated. In longitudinal sections, the CP ended at a zone characterized by the accumulation of electron-dense materials at around the outer microtubule doublets (Figure 2D left, dark arrow). As this zone positionally corresponds to the ‘basal plate’ or ‘axonemal plate’ that is observed in cilia of unicellular organisms and thought to be the site for CP nucleation (Tucker, 1971; Gilula and Satir, 1972; Dute and Kung, 1978; Hoog *et al*., 2014; Dean *et al*., 2016), we also call it ‘basal plate (BP)’, although this structure appears to have a cylindrical shape in the case of airway cells. Closer observations showed that the CP structure began at around the basal end of the BP, but was often sticking out of this BP structure by 20 nm or so (Figure 2D, white arrow). Axonemes in *Camsap3*-mutated cells appeared to have a similar zone, except that microtubules were not always detectable in the BP of mutant samples, as seen in upper axonemal zones (Figure 2D right, dark arrow).

To confirm these observations, we used transverse sections of an airway cell which are slightly tilted, as these allowed us to observe sequential changes in the inner structures of cilium and BB along the longitudinal axis within a single focal plane. In wild-type samples, we detected a group of ciliary sections, each of which exhibits an electron-dense outer ring with a centrally located CP (Figure 2E, left), just as seen in the BP defined above, leading us to judge that this is a transverse view of the BP. In the cilia of *Camsap3*-mutated cells, this structure was deformed showing various abnormalities, such as missing CP (Figure 2E, BP1), overly condensed central materials (Figure 2E, BP2), and partial distortion of the outer ring (Figure 2E, BP3). Some of these BPs retained microtubules in the center (Figure 2E, BP4). Thus, not only the CP but also the entire BP architecture appeared abnormal in the mutant.

These sections also allowed us to examine the TZ, which is characterized by the presence of Y-shaped linkers. We did not find any difference in TZ structures between wild-type and mutant cilia, except that a small number of wild-type TZ contained CP (Figure 2E, BP-TZ); this sample probably represents a section at the boundary region of BP and TZ. The structure of the BBs also looked normal in *Camsap3*-mutated cells, although they tended to lose their rotational polarity in mutant cells, as assessed by measuring the direction of the ‘basal foot’ which normally faces the oral side in wild-type cells (Figure 2F).

### Concentration of CAMSAP3 around the proximal ends of cilia

To begin investigating how CAMSAP3 contributes to microtubule assembly in axonemes, we observed its distribution in airway epithelial cells of adult trachea by immunofluorescence (IF) microscopy, using samples fixed with paraformaldehyde (PFA). The results showed that CAMSAP3 was concentrated around the apical plasma membranes (Figure 3A), as assessed by co-immunostaining for EZRIN, which is known to associate with the apical membranes or microvilli (Berryman *et al*., 1993), as observed in other epithelial cells (Toya *et al*., 2016; Robinson *et al*., 2020; Kimura *et al*., 2021; Usami *et al*., 2021). This position of CAMSAP3 corresponded to the proximal edge of strong α-tubulin IF signals that should represent ciliary microtubules (Figure 3B). Weak CAMSAP3 IF stains were also detectable along cilia as well as in the cytoplasm. We then examined airway epithelia of *Camsap3* mutant mice and found that the mutated CAMSAP3 molecules (CKK domain truncated CAMSAP3) expressed in the mice also accumulated at the apical zones, overlapping with EZRIN (Figure 3A) or ciliary microtubules at their proximal ends (Figure 3B). Curiously, however, the mutated molecule tended to diffuse into upper portions of cilia, even though its expression level was lower than that of the wild-type one (Figure 3C), suggesting that the truncated CAMSAP3 penetrates into the ciliary body more readily than the wild-type CAMSAP3.

**Figure 3.**
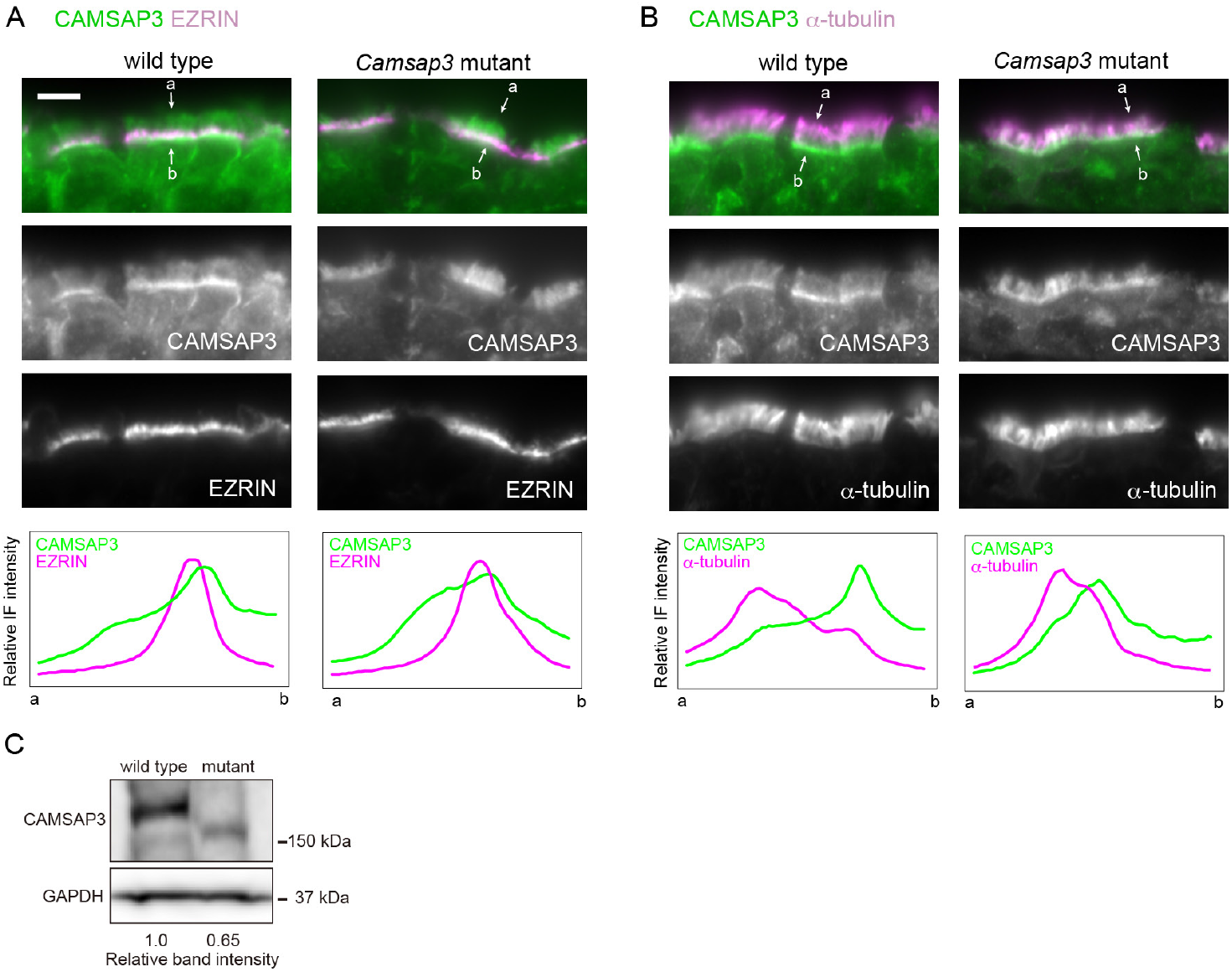
Immunostaining localization of CAMSAP3 in tracheal epithelium. (**A, B**) Trachea derived from P176 mice were fixed with 2% PFA, and processed for co-immunostaining for CAMSAP3 and EZRIN (A) or α-tubulin (B). Images were recorded by conventional immunofluorescence microscopy. Immunofluorescence signals were scanned along the line (not shown) drawn between the two arrows, across the cilia–BB complex from its apical (a) to basal (b) side. A typical image of three to five sections is shown for each set of immunostaining. Scale bars, 10 μm. (**C**) Western blots for CAMSAP3 in trachea isolated from P172 wild-type or *Camsap3*^*dc/dc*^ mice.

### Localization of CAMSAP3 at three distinct sites

To gain more detailed information on CAMSAP3 localization in the cilia and BBs, we co-immunostained CAMSAP3 and components of the cilium-BB complex, analyzing their IF images by Airyscan confocal laser-scanning microscopy, which can yield a 140 nm lateral resolution. For this experiment, we used methanol for fixation of tissues, since some of the proteins to be examined did not give reproducible IF signals in PFA-fixed specimens. The analysis first revealed that CAMSAP3 IF signals, which are condensed at the apical zone of a cell, were split into at least three rows of CAMSAP3 puncta designated #1, #2 and #3 (Figure 4A). The relative IF intensity of the puncta in one row to another row varied from sample to sample; an example of alternative CAMSAP3 patterns is shown in Figure S1. The puncta at the #1 row tended to be variable in morphology, contrasted with a rounded appearance in other rows. Additional puncta were also detectable outside the three rows, but their presence or location was not fixed. Individual CAMSAP3 puncta comprising the three rows were also aligned along the apicobasal axis of the cell, likely due to their association with a single cilium-BB complex at different levels of the apicobasal axis. The distances (mean ± SD) between #1 and #2 and between #2 and #3 along the longitudinal axis were 476 ± 126 nm (n = 30) nm and 364 ± 47 nm (n = 34), respectively.

**Figure 4.**
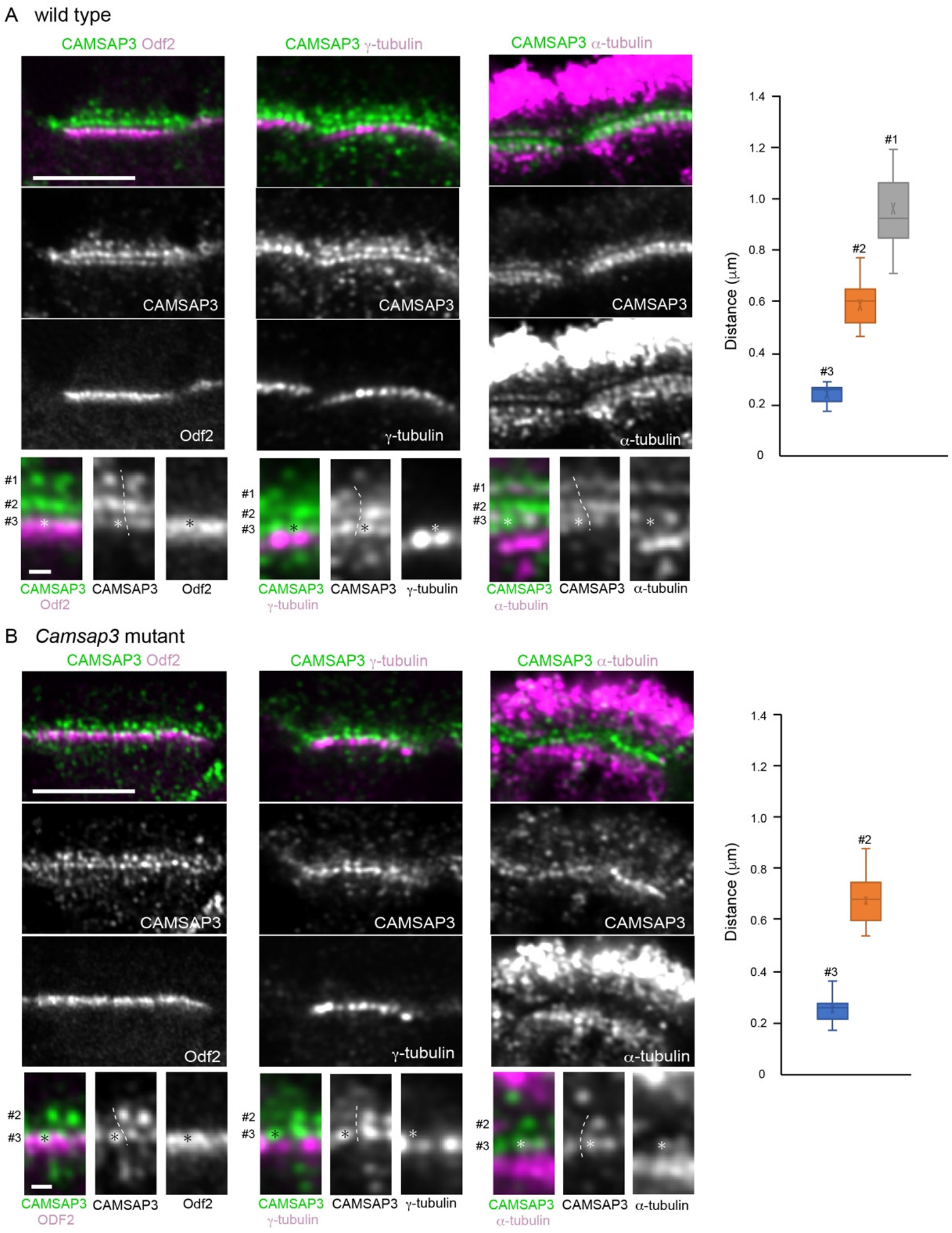
Airyscan microscopic analysis of CAMSAP3 distribution in wild-type and *Camsap3*-mutated multi-ciliated cells. Longitudinal section of multi-ciliated cells in wild-type (**A**) or *Camsap3*-mutated (**B**) trachea, which were co-immunostained for the molecules indicated and recorded by Airyscan microscopy. Part of each image was enlarged at the bottom, in which the numbers indicate three different sites of CAMSAP3 puncta. Asterisks mark an example of CAMSAP3 puncta (or their positions) at the #3 site in each image. Dotted lines indicate a putative boundary between the two sets of CAMSAP3 puncta aligned along the apicobasal axis. A typical image was selected from 8 to 10 sections of tissues, each of which contain several cells that can be analyzed, for each set of immunostaining. The samples were fixed with methanol. The graphs show the distance of CAMSAP3 puncta at different rows from γ-tubulin puncta. P91, P86 and P176 mice were used for immunostaining for Odf2, γ-tubulin, and α-tubulin, respectively. Scale bars, 5 and 0.5 μm in the original and enlarged images, respectively.

To determine the structural relationship between these CAMSAP3 puncta and the BB, we co-immunostained for CAMSAP3 and Odf2 which is known to localize at the transition fibers and basal feet (Kunimoto *et al*., 2012). Odf2 overlapped with the #3 CAMSAP3 puncta at the upper portion of the former, further extending down to CAMSAP3-negative zones (Figure 4A, leftmost). γ-Tubulin, which has been reported to accumulate at the basal foot as well as at additional lower regions of BB in motile cilia (Hagiwara *et al*., 2000; Clare *et al*., 2014), was detected below the #3 puncta, occasionally overlapping with faint CAMSAP3 IF signals located underneath the #3 row (Figure 4A, middle). The distances (mean ± SD) between γ-tubulin and the #1, #2 or #3 CAMSAP3 were estimated at 962 ± 130 (n = 25), 592 ± 89 nm (n = 18) and 240 ± 37 nm (n = 19), respectively (Figure 4A graph).

We then detected microtubules by α-tubulin immunostaining (Figure 4A, rightmost). Cytoplasmic microtubules formed meshwork below the #3 CAMSAP3 puncta, and some of them entered into the #3 zone, yielding their overlapping view. On the other hand, ciliary microtubules, which should have a structural continuity to the basal body, were not clearly detectable at their proximal regions in our methanol-fixed samples for unknown reasons. Nevertheless, CAMSAP3 puncta at the #1 site exhibited some overlapping with microtubule clusters.

We next analyzed the distribution of the CKK domain-truncated CAMSAP3 by Airyscan microscopy in *Camsap3*-mutated cells (Figure 4B). As seen for wild-type CAMSAP3, the mutant molecules formed multiple rows. Co-immunostaining for CAMSAP3 and Odf2, γ-tubulin or α-tubulin showed that their spatial relationships were similar to those found in wild-type cells. However, we could not convincingly detect the #1 row, although CAMSAP3 IF signals were randomly present at the ciliary zone (Figure 4B). We also noted that about half of CAMSAP3 puncta in the #2 row were missing. These suggest that, although the mutated CAMSAP3 still associates with the cilium-BB system at the #3 site, it does not do so at the #1 site, and its binding to the #2 site also became unstable. The distance (mean ± SD) between the #2 and #3 puncta was 335 ± 46 nm (n = 23), and those between γ-tubulin and the #2 or #3 puncta were 674 ± 96 (n = 17) and 250 ± 49 nm (n = 17), respectively (Figure 4B graph).

### Localization of CAMSAP3 puncta at the transition zone

For more precise localization of CAMSAP3 puncta, we co-immunostained wild-type ciliated cells for CAMSAP3 and Chibby, a protein that localizes at a proximal region of TZ, playing a role in ciliogenesis (Enjolras *et al*., 2012; Burke *et al*., 2014). Confocal microscopy of longitudinal sections showed that Chibby localizes above the #3 CAMSAP3 puncta (Figure 5A). We then used Airyscan microscopy, confirming that major Chibby puncta are located above the #3 puncta. To relate the distribution of the #2 and #3 CAMSAP3 puncta with the actual structure of the cilia-BB complex, we overlaid their positions on its ultrastructural image, referring to the published location of γ-tubulin (Clare *et al*., 2014), Odf2 (Kunimoto *et al*., 2012) and Chibby (Burke *et al*., 2014) in motile cilia, which were determined at the ultrastructural levels (Figure 5B). This comparison of IF images with the ultrastructure suggests that the #2 CAMSAP3 puncta are localized at a distal portion of TZ or at BP, while the #3 CAMSAP3 puncta, at an upper region of the BB.

**Figure 5.**
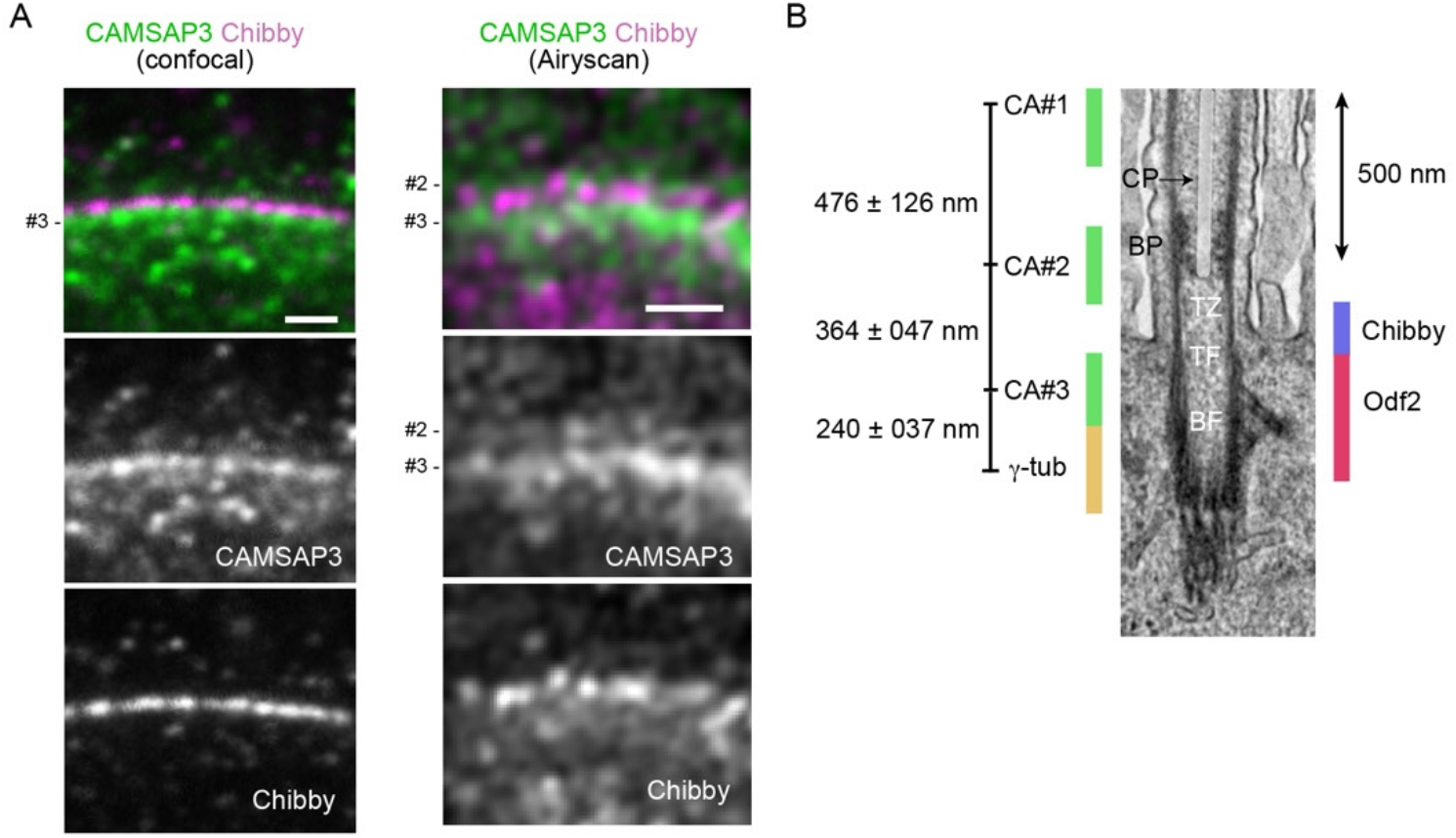
CAMASP3 localization in the transition zone. **(A)** Longitudinal section of a wild-type multiciliated cell, co-immunostained for CAMSAP3 and Chibby, and recorded by confocal or Airyscan microscopy. In these samples, the relative IF intensity of #3 CAMSAP3 puncta is high. Trachea were collected from P180 mice and fixed with methanol. Scale bar, 1 μm. **(B)** A putative map of CAMSAP3 puncta in relation to the ultrastructure of the axoneme, transition zone and basal body. The relative position of each CAMSAP3 punctum was determined by measuring the distances between the puncta, and then the position of the entire CAMSAP3 puncta relative to the ultrastructure was adjusted by placing the #3 punctum at the level of the upper half of Odf2 distribution, as observed in Figure 4. The positions of Chibby, Odf2 and γ-tubulin were estimated referring to previous publications, although γ-tubulin localization needs further confirmation. The scale of the ultrastructure image is shown by a 500 nm bar. The CP is overlaid with a semitransparent white color. CP, the central pair of microtubules; BP, basal plate; TZ, transition zone; TF, transition fiber; BF, basal foot; γ-tub, γ-tubulin; CA, CAMSAP3.

Finally, we viewed the overall distribution pattern of basal bodies and microtubules from the top of ciliated cells in which GFP-tagged Centrin 2, a basal body protein, is genetically expressed (Higginbotham *et al*., 2004). Basal bodies exhibited typical horizontal arrays in wild-type cells, and a similar pattern was maintained in *Camsap3*-mutated cells (Figure S2). Microtubules form a horizontal network at the inter-BB spaces (Kunimoto *et al*., 2012; Tateishi *et al*., 2013; Tateishi *et al*., 2017), and we did not find any particular change of this network in *Camsap3*-mutated cells (Figure S2, and Videos 5 and 6). Microtubules located below the basal bodies also formed networks, and their pattern was indistinguishable between wild-type and *Camsap3*-mutated cells. Thus, it seems that *Camsap3* mutation did not affect the overall BB assembly, nor the horizontal networks of microtubules associated with the BBs.

## Discussion

The present study demonstrated that CAMSAP3 concentrates at multiple sites of the cilium-BB complex in tracheal motile cilia in mice, and dysfunction of this molecule results in unsynchronized beating of multicilia, as well as the loss of CP and other structural defects in this complex. Comparison of CAMSAP3 distribution with the ultrastructure of a cilium suggested that the #2 site corresponds to BP or an upper region of TZ, and the #1 and 3 sites are located more distal and proximal than the BP, respectively. A preceding study using nasal epithelial cells also reported that a *Camsap3* hypomorphic mutation caused similar functional and structural defects in the multicilia (Robinson *et al*., 2020). This study, however, detected CAMSAP3 as a single dot located between the axoneme and BB in P30 mice, contrasted with our observation that the CAMSAP3-bearing region is subdivided into multiple sites. Two possibilities can be considered to explain the difference between the two observations. First, CAMSAP3 may interact with ciliary components in different ways between the nasal and tracheal multiciliated cells, and secondly, different sample preparation or microscopic methods can lead to different results. Actually, we noted a variation in the CAMSAP3 distribution pattern. Whichever the case is, the CAMSAP3 dot in P30 nasal cilia may correspond to the #2 or #3 puncta in tracheal cilia, considering their distance from γ-tubulin of the BB. In nasal motile cilia, an additional dot overlapping with γ-tubulin was also observed in P3 mice (Robinson *et al*., 2020), but we did not examine tracheal motile cilia in such young mice.

The information that CAMSAP3 localizes around at the BP enabled us to discuss its potential role in CP formation on the basis of our knowledge about how CAMSAP3 controls microtubules. That is, it stabilizes the minus-end of a microtubule and simultaneously serves to tether the microtubule filament to particular subcellular sites (Nashchekin *et al*., 2016; Toya *et al*., 2016). Here, we propose that CAMSAP3 may function to stabilize the CP by binding to the minus end of microtubules at around the BP, as the minus ends of axonemal microtubules are oriented basally (Euteneuer and Mcintosh, 1981). Removal of this mechanism would have led to a collapse of the CP. Notably, the CKK-truncated CAMSAP3 expressed in *Camsap3*^*dc/dc*^ mice, which is unable to bind the minus-ends of microtubules, were still detected at the #2 site but not always. It is possible that CAMSAP3 can stay at this site without binding microtubules, but its association with a putative binding partner became unstable under this situation, as observed in previous studies (Toya *et al*., 2016), and consistent with the idea that CAMSAP3 does interact with microtubules at this site. To test these ideas, however, it will be prerequisite to determine whether CAMSAP3 is located inside or outside the axoneme for its interaction with microtubules.

Since BP architecture was also perturbed in *Camsap3*-mutated airway cells, the presence of CAMSAP3 or CP may also play a role in maintaining this structure. On the other hand, detection of CAMSAP3 puncta at regions more distal than BP (at the #1 site) suggest that CAMSAP3 may interact with microtubules at the axoneme, because its puncta disappeared in *Camsap3*-mutated cilia. This could be another site for CAMSAP3 to support the CP, but its potential function remains to be further investigated. Mutated CAMSAP3 tended to diffuse into the upper regions of a cilium. This might have occurred due to the release of CAMSAP3 from the #1 or #2 sites.

Loss of CP may explain why the *Camsap3*-mutated motile cilia showed unsynchronized beating, since CP deficiency due to various mechanisms is known to coincide with disorganized ciliary movement (Wilson *et al*., 2009; Kott *et al*., 2013; Nozawa *et al*., 2013; Zhu *et al*., 2019). Defective CP may also have been involved in the perturbation of BB polarity observed in the absence of functional CAMSAP3. For example, the aberrant beating of multicilia in *Camsap3*-mutated cells cannot produce sufficient hydrodynamic forces that are required for the polarized alignment of them (Guirao *et al*., 2010). On the other hand, we cannot exclude the possibility that CAMSAP3 associated with the cilia-BB complex could have some direct role in determining BB polarity. Defects in this putative mechanism might have perturbed BB polarity and in turn induced uncoordinated beating of cilia.

What is the function of CAMSAP3 at the #3 site that corresponds to the distal part of the BB? It overlaps with the upper component of Odf2 that is known to associate with transition fibers as well as basal feet, removal of which results in loss of BB but without affecting CP (Kunimoto *et al*., 2012). Despite their overlapping, CAMSAP3 mutants did not mimic what is observed in Odf2 mutants, suggesting that they are not functionally linked to one another. The loss of CAMSAP3 function did not affect the horizontal network of microtubules interspaced between BBs, suggesting that their assembly of microtubes is regulated by factors other than CAMSAP3, such as γ-tubulin, an authentic microtubule nucleator. Thus, the role of CAMSAP3 at the BB remains mysterious. It is noted that the CKK-deficient CAMSAP3 puncta were normally detectable at the #3 site, implying that CAMSAP3 proteins cluster here independently of microtubules.

Curiously, the role of CAMSAP3 in multiciliated cells appears not thoroughly conserved among different organs, contrasted with many other ciliary components whose loss induces PCD. In multiciliated ependymal cells of the brain, CAMSAP3 does not particularly associate with the cilium-BB complex, and their ciliary beating occurs normally in *Camsap3*-mutant mice (Kimura *et al*., 2021). In the ependymal cells, CAMSAP3 localizes under the apical plasma membranes and its major role is to regulate the organization of cortical microtubules that are required for establishing a microtubule-dependent signaling system. These observations suggest that CAMSAP3 contributes to multiciliary organization in a tissue-specific manner, although the possibility that other CAMSAP subtypes or unidentified similar proteins have taken over the role of CAMSAP3 in ependymal cells is not excluded. It should also be noted that the CAMSAP family or CKK domain-containing-proteins have been detected only in the metazoan (Baines *et al*., 2009), which implies that their role in the ciliary system is limited to particular animal species. Future studies will be necessary to reveal how CAMSAP3 becomes associated with the cilium–BB complex and supports its structure and function in particular tissues and species.

## Materials and Methods

### Mice

A *Camsap3* mutant mouse line, *Camsap3*^*dc/dc*^, was described previously (Toya *et al*., 2016). The transgenic mouse line expressing GFP-Centrin2 (EGFP-CETN2) was a gift from Holden Higginbotham, Brigham Young University (Higginbotham *et al*., 2004). Female mice of this transgenic line were crossed with *Camsap3*^*dc/+*^ males to generate those expressing GFP-Centrin2 under *Camsap3* mutation. ICR mice (Slc; ICR, Japan SLC) were used to for immunostaining analysis of Chibby. The age of mice used for the study was recorded for each experiment and described in the legends of figures. The experiments using mice were performed in accordance with the protocol(s) approved by the Institutional Animal Care and Use Committee of the RIKEN Center for Biosystems Dynamics Research (RIKEN Kobe Branch), or the Guidelines of Animal Experimentation of National Institutes for Natural Sciences.

### Scanning electron microscopy

Mice were anesthetized with isoflurane, and dissected to reveal trachea and cut diaphragm. The tracheas were prefixed with a fixative solution (2% paraformaldehyde and 2.5% glutaraldehyde in 0.1M cacodylate buffer, pH 7.4) for 5 min, then removed from mice. The trachea was cut into small pieces, and the samples were immersed in the same fixative solution at room temperature for 2 hr. The samples were washed with 0.1M cacodylate buffer and postfixed in 1% OsO4 in 0.1M cacodylate buffer on ice for 2 hr. Then the samples were washed with distilled water, and stained overnight with 0.5% aqueous uranyl acetate at room temperature. The stained samples were dehydrated with ethanol the samples were rinsed with distilled water, and further stained with 0.5% uranyl acetate solution overnight at room temperature. The samples were rinsed with distilled water and dehydrated through a graded series of ethanol, transferred to isopentyl acetate and critical point-dried using liquid CO_2_. The dried specimens were evaporation-coated with Osmium and examined in a JSM-5600LV scanning electron microscope at an accelerating voltage of 10 kV.

### Transmission electron microscopy

Tracheas were fixed with 2% paraformaldehyde and 2.5% glutaraldehyde in 0.1M cacodylate buffer (pH 7.4) at 4°C overnight. After washing five times with 0.1M cacodylate buffer, the specimens were further fixed with 1% OsO4 in 0.1M cacodylate buffer at 4°C for 1 h and then washed five times with Milli-Q water. The specimens were incubated in 1% uranyl acetate overnight and washed five times with Milli-Q water. Specimens were dehydrated with ethanol series (20%, 50%, 70%, 90% and 99.5% for 5 min each, and twice in 100% for 10 min) and acetone, then followed by infiltration with Epon 812 (TAAB). Polymerization was performed at 60°C for 72 h. Samples were cut with an ultramicrotome (ULTRA CUT UCT, Leica) equipped with a diamond knife (Diatome), doubly stained with uranyl acetate and lead citrate, and examined in a transmission electron microscope (JEM-1400 Plus ; JEOL) operated at 100 kV.

### Immunofluorescence staining and microscopy

Isolated trachea tissue was fixed in a PEM buffer (PEM: 100 mM PIPES, 5 mM EGTA, and 5 mM MgCl_2_, adjusted to pH 6.9 with 2 M KOH) containing 2% paraformaldehyde and 0.1 M sorbitol for 1 h at room temperature, except that the tissue was treated with 4% paraformaldehyde at 37°C for 1 hr and then at 4°C for 24 hr for microtubule fixation. Alternatively, trachea tissue was fixed with methanol for 20 to 45 min at – 20°C, followed by treatment with acetone for 1 min at –20°C. After fixation, the tissue was rinsed three times with PEM for 10 min. Then, cryoprotection was carried out by incubation with 15% sucrose in PEM overnight at 4°C, followed by incubation with 20% sucrose in PEM for 6 h, and 30% sucrose in PEM overnight at 4°C. The tissue was frozen in OCT compound in an embedding mold (Polysciences, Inc.) in liquid nitrogen and stored at –80°C. Frozen samples were sliced at 7 mm thickness with a cryostat (HM500M, MICROM).

After washing twice with PBS, the sections were blocked with Blocking One (Nacalai Tesque) for 1 hr at room temperature, followed by incubation with primary antibodies in Blocking One (Nacalai Tesque) overnight at 4°C. Sections were washed two times with PBS and incubated for one hour at RT with corresponding secondary antibodies. Stained sections were washed two times with PBS and mounted with either Fluorsave (Chemicon) or 2,2′-thiodiethanol (Staudt *et al*., 2007). For samples used for Airyscan microscopy, antigen retrieval was performed by incubating sections in HistoVT One (Nacalai Tesque) for 3 min at 95°C, prior to the incubation with primary antibodies.

For conventional immunofluorescence microscopy, samples were analyzed by Zeiss Axiopan 2 through Plan-APOCHROMAT 63x/1.4 Oil Dic or Plan-NEOFLUAR 40x/1.3 Oil Dic objectives. For confocal microscopy, a NIKON A1 microscope with Nikon Plan Apo Vc 100x/1.40 oil OFN25 DIC N2 objective lens was used. For super-resolution microscopy, stacks of images were taken along the z-axis at optimal intervals using Airyscan (LSM880, Zeiss) with 63x/1.40 NA and a 100x/1.46 NA objective lenses. TCS SP8 STED with HCX PL APO CS 100x/1.40 oil immersion objective lens (Leica) was used to acquire the horizontal views of cells, and the optical conditions for this imaging are shown in Supplementary Table 1. Acquired images were processed using Zen (Zeiss), Adobe Photoshop (Adobe), or ImageJ (Wayne Rasband, NIH).

### Measurement of distances between CAMSAP3 puncta

To determine the distance between two IF puncta, we measured from the center of one punctum to that of the other punctum, using ImageJ. Three to ten sections were used for each measurement.

### Quantification of ciliary orientation in multiciliated epithelial cells

To quantify the alignment of multicilia along the longitudinal axis of trachea, the angles of cilia in SEM images were measured using ImageJ. Briefly, a line was drawn along individual cilia and then the angle of the line was measured. Measured angles of cilia were normalised by the average angle of all the cilia within the cell in order to eliminate a possible difference in the orientation of multiciliated epithelial cells. Polar plots for the distribution of ciliary angles were generated by ggplot2 in R. For the statistical test, we applied the F-test function (var.test) in R.

### Live imaging of ciliary motion and quantification of data

Live images of cilia were acquired as described previously (Shinohara *et al*., 2015). Briefly, tracheas were collected into HEPES-buffered Dulbecco’s modified Eagle’s medium (pH7.2) supplemented with 10% fetal bovine serum. Motility of cilia was examined at 25°C with a high-speed CMOS camera (HAS-U2, DITECT) that was connected to a microscope (BX53, Olympus) with a 40x objective lens (UPLFLN40X, Olympus) at a frame rate of 300 frames/s for tracheal cilia. To emphasize the movement of cilia, time-series images were adjusted to gain proper contrast and brightness after subtracting the minimum value through the images from each frame using ImageJ.

The synchrony of the motion of the cilia was quantified by calculating the flow field of the cilia carpet by PIV analysis. We denoised the time-lapse images (128 pix x 128 pix) by a Gaussian filter followed by the enhancement of the contrast using Fiji plugins. Then, the flow field of the time-lapse images was obtained by PIV analysis using a python library (OpenPIV). The interrogation window of PIV was 32 pix, and the overlap between adjacent interrogation areas was 24 pix.

### Antibodies

The following primary antibodies were used: Rabbit anti-CAMSAP3 (Tanaka et al., 2012, 1: 100), mouse anti-α-tubulin (clone DM1A, Sigma-Aldrich T9026, 1:1000), rabbit monoclonal anti-β-tubulin (clone 9F3, Cell Signaling Technology Cat#2128, 1: 100), mouse anti-γ-tubulin (Sigma-Aldrich T6557, 1:1000), rabbit anti-γ-tubulin (Sigma-Aldrich T3559, 1:500), mouse anti-EZRIN (Abcam 4069, 1:100), rat anti-Odf2 (a gift from Sachiko Tsukita, Osaka University, 1: 2), mouse monoclonal anti-Chibby (Santa Cruz sc-101551, 1:100). The following secondary antibodies were used: Donkey CF-488A anti-rat IgG (Biotium 20027, 1:000) and CF-568 anti-rabbit IgG (Biotium 20698, 1:1000) antibodies; goat Alexa Fluor-488 anti-rat IgG (Invitrogen, A-11006, 1:500), Alexa Fluor-488 anti-rabbit IgG (Invitrogen, A-11034, 1:500), Alexa Fluor-568 anti-mouse IgG (Invitrogen, A-11031, 1:500), Alexa Fluor-594 anti-rabbit IgG (Invitrogen A-11037, 1:500), Alexa Fluor-647 anti-rabbit IgG (Invitrogen, A-21245, 1:500) and Alexa Fluor-647 anti-mouse IgG (Invitrogen, A-21236, 1:1000) antibodies; and goat STAR580 anti-mouse IgG (Abberior 2-0002-005-1, 1:500) and STAR635P anti-rat IgG (Abberior 2-0132-007-5, 1:500).

### Western blot analysis

Trachea tissues were homogenized in TMEN buffer (20 mM Tris-HCl pH 7.4, 1 mM MgCl_2_, 2 mM EDTA, and 150 mM NaCl) containing the proteinase inhibitor cocktail cOmplete ultra (Roche, 5892970001) and the phosphatase inhibitor cocktail PhosSTOP (Roche, 4906845001). Then, 2% (w/v) SDS solution was added to homogenized trachea tissues. Proteins were separated by 4-20% gradient SDS-PAGE gel (Cosmobio, 414879) and then transferred to PVDF membrane (Millipore, IPVH00010). To detect CAMSAP3 and GAPDH for a loading control, anti-CAMSAP3 rabbit antibody (1:1,000) and anti-GAPDH mouse antibody (MBL, M171-3, 1:1,000) were used as primary antibodies. Quantification of band intensities was done using ImageJ.

### Audio recording

Respiratory sound of mice was recorded using the Voice Memos application installed in iPhone XS (Apple).

## Acknowledgements

We thank Hiroshi Hamada for his support of live imaging experiments, and Mitsuru Morimoto for his advice on the study of lung, Sachiko Tsukita for anti-Odf2 antibody, and Ritsu Kamiya for critical reading of the manuscript. This work was supported by the program Grant-in-Aid for Scientific Research (S) (grant number 25221104) from the Japan Society for Promotion of Science to M.Ta; the Japan Society for the Promotion of Science, KAKENHI (grant numbers 17H03689 and 16H06280) and the Japan Science and Technology Agency, Core Research for Evolutional Science and Technology (JPMJCR1654) to T.F.; the Japan Society for the Promotion of Science, KAKENHI (grant number JP20K06645) to M.To; and Daiichi Sankyo Foundation of Life Science to M.S.

## Author contributions

Conceptualization: M.Ta., M.To.; Formal analysis: T.Y., T.S., Y.K.; Investigation: H.S., F.M-U., T.K., T.I., K.O., S.O., K.M., Y.S., Y.K., M.To.; Writing-original draft: M.Ta.; Writing-reviews & editing: M.To., T.F., F.M-U.; Supervision: M.Ta., T.F., S.Y., M.S.; Project administration: M.Ta.; Funding acquisition: M.Ta., T.F.

---

**Supplementary Figure S1.**
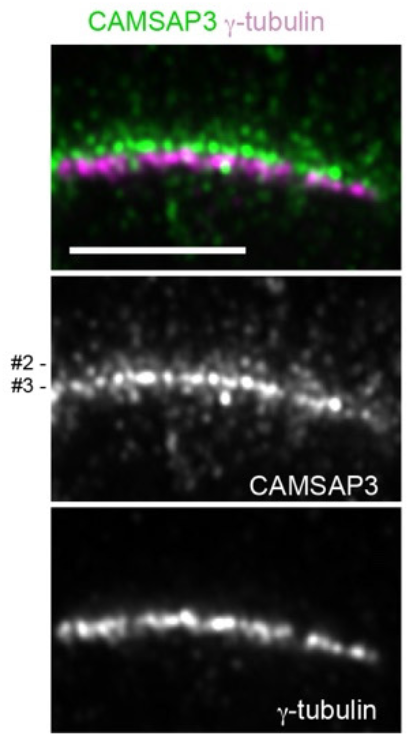
An alternative pattern of CAMSAP3 distribution. Longitudinal section of a multi-ciliated cell in wild-type trachea, which was co-immunostained for CAMSAP3 and γ-tubulin and recorded by Airyscan microscopy. Note that the #3 row, identified by its position next to γ-tubulin, shows the most intense IF signals in this particular specimen.

**Supplementary Figure S2.**
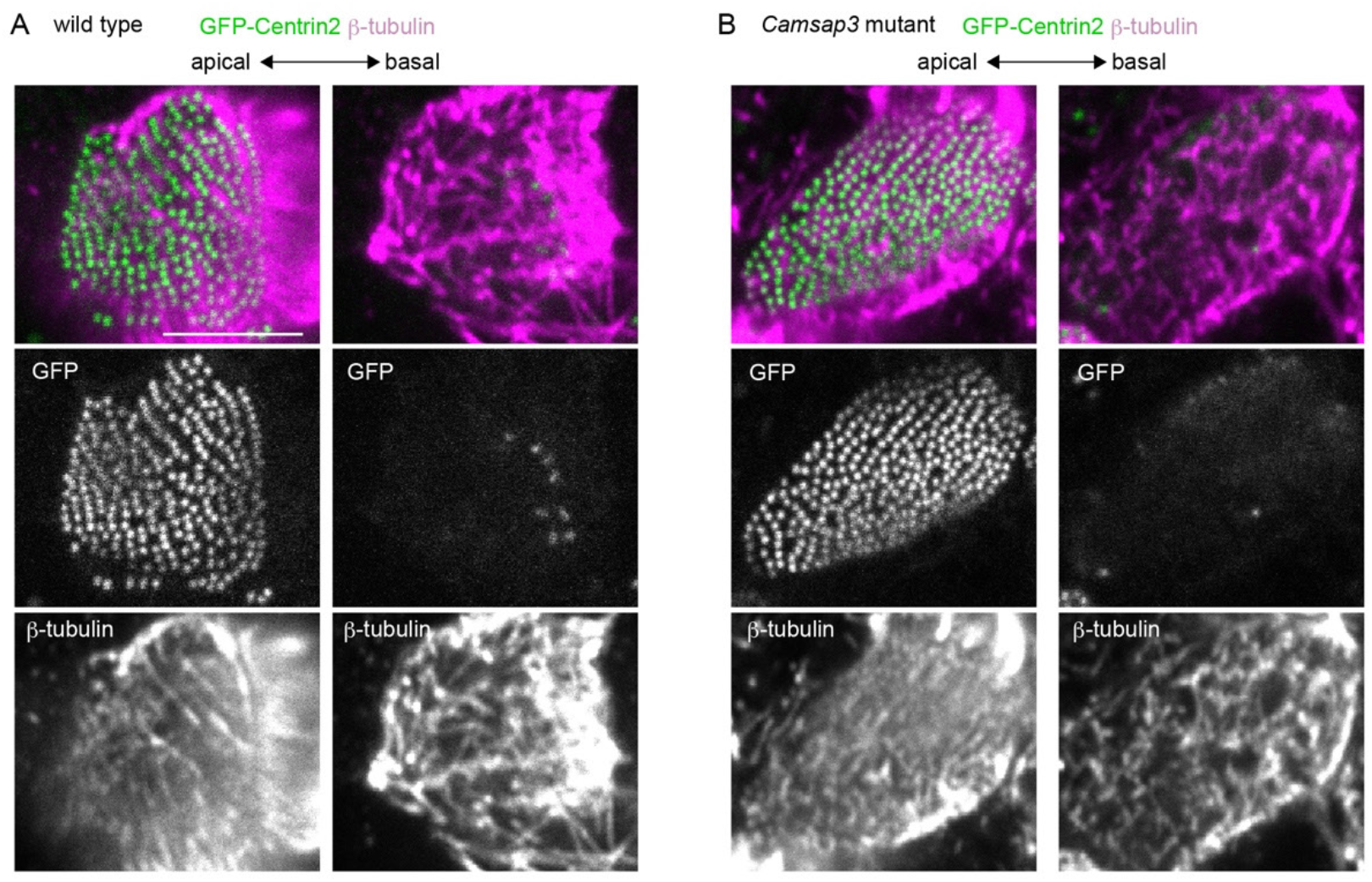
Microtubule distribution at the apical portions of a multiciliated cell. β-tubulin was immunostained using trachea isolated from a wild-type (A) or *Camsap3*^*dc/dc*^ (B) mouse expressing GFP-Centrin2. The images at two horizontal levels, which are 0.12 (in A) and 0.10 μm (in B) apart, are shown. Oral side is at the right. P180 mice were used. The same samples were used in Videos 5 and 6. Scale bars, 5 μm.

## Video Legends

**Video 1**. Beating multicilia in a wild-type airway epithelial cell. The time-lapse images were acquired at 300 fps and displayed at 31 fps.

**Video 2**. Beating multicilia in a *Camsap3*-mutated airway epithelial cell. The time-lapse images were acquired at 300 fps and displayed at 31 fps.

**Video 3**. Beating multicilia in a wild-type airway epithelial cell, which is overlaid with a flow vector field calculated by PIV analysis. The flow vector field was also shown separately at the right. The scale arrow and scale bar are 20 μm/s and 1 μm, respectively. Part of Video 1 was used for this analysis.

**Video 4**. Beating multicilia in a *Camsap3*-mutated airway epithelial cell, which is overlaid with a flow vector field calculated by PIV analysis. The flow vector field was also shown separately at the right. The scale arrow and scale bar are 20 μm/s and 1 μm, respectively. Part of Video 2 was used for this analysis.

**Video 5**. Animation of sequential optical sections, each of which is 0.2 μm thick, of a wild-type multi-ciliated epithelial cell, in which GFP-Centrin (green) and β-tubulin (magenta) are visualized by fluorescence signals. The animation view begins around the level where the array of GFP-Centrin is detectable, then shifts toward a more basal view of the cell at a speed of 2 fps. See also Figure 6A.

**Video 6**. Animation of sequential optical sections, each of which is 0.2 μm thick, of a *Camsap3*-mutated multi-ciliated epithelial cell, in which GFP-Centrin (green) and β-tubulin (magenta) are visualized by fluorescence signals. The animation view begins at the level where the array of GFP-Centrin is detectable, then shifts toward a more basal view of the cell at a speed of 2 fps. See also Figure 6B.

## Audio Legends

**Audio 1**. Respiratory sound of a P50 *Camsap3* mutant mouse.

